# Broad-spectrum virucidal activity of bacterial secreted lipases against flaviviruses, SARS-CoV-2 and other enveloped viruses

**DOI:** 10.1101/2020.05.22.109900

**Authors:** Xi Yu, Liming Zhang, Liangqin Tong, Nana Zhang, Han Wang, Yun Yang, Mingyu Shi, Xiaoping Xiao, Yibin Zhu, Penghua Wang, Qiang Ding, Linqi Zhang, Chengfeng Qin, Gong Cheng

**Author notes:** Address correspondence to: Gong Cheng Ph.D., School of Medicine, Tsinghua University, Beijing, P.R. China, 100084. These authors contributed equally to this work (Xi Yu, Liming Zhang, Liangqin Tong).

## Abstract

Viruses are the major aetiological agents of acute and chronic severe human diseases that place a tremendous burden on global public health and economy; however, for most viruses, effective prophylactics and therapeutics are lacking, in particular, broad-spectrum antiviral agents. Herein, we identified 2 secreted bacterial lipases from a *Chromobacterium* bacterium, named *Chromobacterium* antiviral effector-1 (*Cb*AE-1) and *Cb*AE-2, with a broad-spectrum virucidal activity against dengue virus (DENV), Zika virus (ZIKV), severe acute respiratory syndrome coronavirus 2 (SARS-CoV-2), human immunodeficiency virus (HIV) and herpes simplex virus (HSV). The *Cb*AEs potently blocked viral infection in the extracellular milieu through their lipase activity. Mechanistic studies showed that this lipase activity directly disrupted the viral envelope structure, thus inactivating infectivity. A mutation of *Cb*AE-1 in its lipase motif fully abrogated the virucidal ability. Furthermore, *Cb*AE-2 presented low toxicity *in vivo* and *in vitro*, highlighting its potential as a broad-spectrum antiviral drug.

## Main Text

Emerging and re-emerging viral diseases cause a major burden on global public health. Coronavirus disease 2019 (COVID-19), a new severe acute respiratory disease caused by severe acute respiratory syndrome coronavirus 2 (SARS-CoV-2), emerged at the end of December 2019 and is now pandemic globally (Rothan and Byrareddy, 2020; Zheng, 2020). As of May 9^th^, 2020, World Health Organization (WHO) has recorded 3,855,788 confirmed cases and 265,862 deaths across over 200 countries (WHO, 2020). In addition, several highly pathogenic viruses, such as human immunodeficiency virus (HIV), dengue virus (DENV), severe acute respiratory syndrome coronavirus (SARS-CoV), Zika virus (ZIKV) and Ebola virus (EBOV), suddenly emerged/re-emerged and disseminated rapidly (Cordova-Villalobos et al., 2017; Holt, 2018; Obando-Pacheco et al., 2018; Oidtman et al., 2019). There are billions of reported cases of viral infections annually, in which millions of patients suffer from severe clinical symptoms and even death (Peteranderl et al., 2016; Sejvar, 2016; Song et al., 2017; Stein et al., 2017). However, neither specific antiviral drugs nor vaccines will be immediately available when a new virus emerges. Therefore, broad-spectrum antiviral agents are urgently needed to control an emerging crisis of public health (Boldescu et al., 2017; Totura and Bavari, 2019; Zhu et al., 2015), thus reinforcing the arsenal of our antiviral options. Herein, we identify 2 secreted bacterial proteins from a *Chromobacterium* bacterium that can robustly resist infections of DENV, ZIKV, SARS-CoV-2, HIV and herpes simplex virus-1 (HSV-1), suggesting their potential as broad-spectrum antiviral agents.

*Chromobacterium spp.* constitute a genus of gram-negative bacteria that are occasionally pathogenic to humans and animals. A soil bacterium named *Chromobacterium sp.* Panama (*Csp_P*) identified from the midgut of a field-caught *Aedes aegypti* showed strong virucidal activity against DENV (Caragata et al., 2020; Ramirez et al., 2014; Saraiva et al., 2018). In this study, we first identified a *Chromobacterium* bacterium from the gut of *A. aegypti* mosquitoes reared in our insectary facility. We thus refer to this bacterium as *Chromobacterium sp.* Beijing (*Csp_BJ*). We sequenced the whole bacterial genome to characterize *Csp_BJ* and uploaded the whole genome information into the NCBI database (NCBI Taxonomy ID: 2735795). Based on genomic comparison, *Csp_BJ* shares either 99.48% identity to *Chromobacterium haemolyticum CH06-BL* or 99.42% identity to *Chromobacterium rhizoryzae JP2-74* strain. Oral supplementation of this bacteria in *A. aegypti* largely impaired mosquito permissiveness to DENV (Supplementary Figure 1A and 1B) and ZIKV (Supplementary Figure 1C and 1D), suggesting a close relationship between the identified *Csp_BJ* and *Csp_P* strains. We next aimed to understand how *Csp_BJ* resists viral infection in mosquitoes. Bacteria usually exploit many effectors, such as cellular components, metabolites or secreted proteins, to regulate their host immune or physiological status for effective colonization. We therefore identified the bacterial effector(s) that modulate infection of DENV and ZIKV through differential fragmentation. In this experiment, we cultured *Csp_BJ* for 24 hr at 30°C. The cell-free culture supernatant was collected by centrifugation and filtration through a 0.22 µm filter unit, whereas the cell lysates were generated by sonication. Either the bacterial cell lysate or the culture supernatant was mixed with 50 plaque-forming units (pfu) of DENV or ZIKV and incubated for 1 hr, and then the infectious viral particles were determined by a plaque forming assay (Figure 1A). Incubation of the culture supernatant but not the bacterial lysates resulted in significant suppression of DENV (Figure 1B) and ZIKV (Figure 1C) infectivity in Vero cells, indicating that an extracellular effector(s) secreted by *Csp_BJ* was responsible for viral inhibition. Next, we investigated whether the effector(s) was a secreted protein, small peptide, lipid, polysaccharide or other metabolite. Therefore, the culture supernatant was separated using a 3 kDa-cutoff filter (Wu et al., 2019). Either the upper retentate (proteins and large peptides) or the lower liquid filtrate (small molecule compounds and short peptides) was mixed with the viruses for incubation in Vero cells (Figure 1D). Intriguingly, inoculation of the retentate rather than the filtrate inhibited the infectivity of DENV and ZIKV (Figure 1E and 1F), suggesting that the effector(s) might be a protein(s) secreted by *Csp_BJ*. Subsequently, the protein components in the upper retentate were separated by SDS-PAGE and then identified by mass spectrometry (Figure 2A). The highly abundant proteins with secretable properties were selected, expressed and purified in an *Escherichia coli* expression system (Figure 2B). Of all the proteins tested, the bacterial protein encoded by gene3771 (Accession: MT473992) significantly impaired DENV and ZIKV infection in Vero cells (Figure 2C and 2D). We named this protein *Chromobacterium* antiviral effector-1 (*Cb*AE-1). Intriguingly, a *Cb*AE-1 homologue with 55.61% amino acid identity named *Cb*AE-2 (Accession: MT473995) was further identified from *Csp_BJ* based on sequence comparison (Supplementary Figure 2A and 2B). Both effectors encoded a lipase domain with a typical GDSL motif (Casas-Godoy et al., 2018). To further confirm the virucidal activity, both proteins were expressed and purified in *E. coli* (Figure 2E). A serial concentration of recombinant proteins was mixed with 50 pfu of DENV or ZIKV for plaque assay in Vero cells. The half-inhibitory concentration (IC_50_) of *Cb*AE-1 was 0.0016 µg/ml for DENV and 0.0026 µg/ml for ZIKV (Figure 2F and 2G). However, the IC_50_ of *Cb*AE-2 for both flaviviruses was 370-2400 times higher than that of *Cb*AE-1, indicating a much more robust virucidal activity of *Cb*AE-1 against flaviviruses (Figure 2F and 2G). Thus, we identified 2 bacterial effectors with a high virucidal activity from a *Csp_BJ* bacterium.

**Figure 1.**
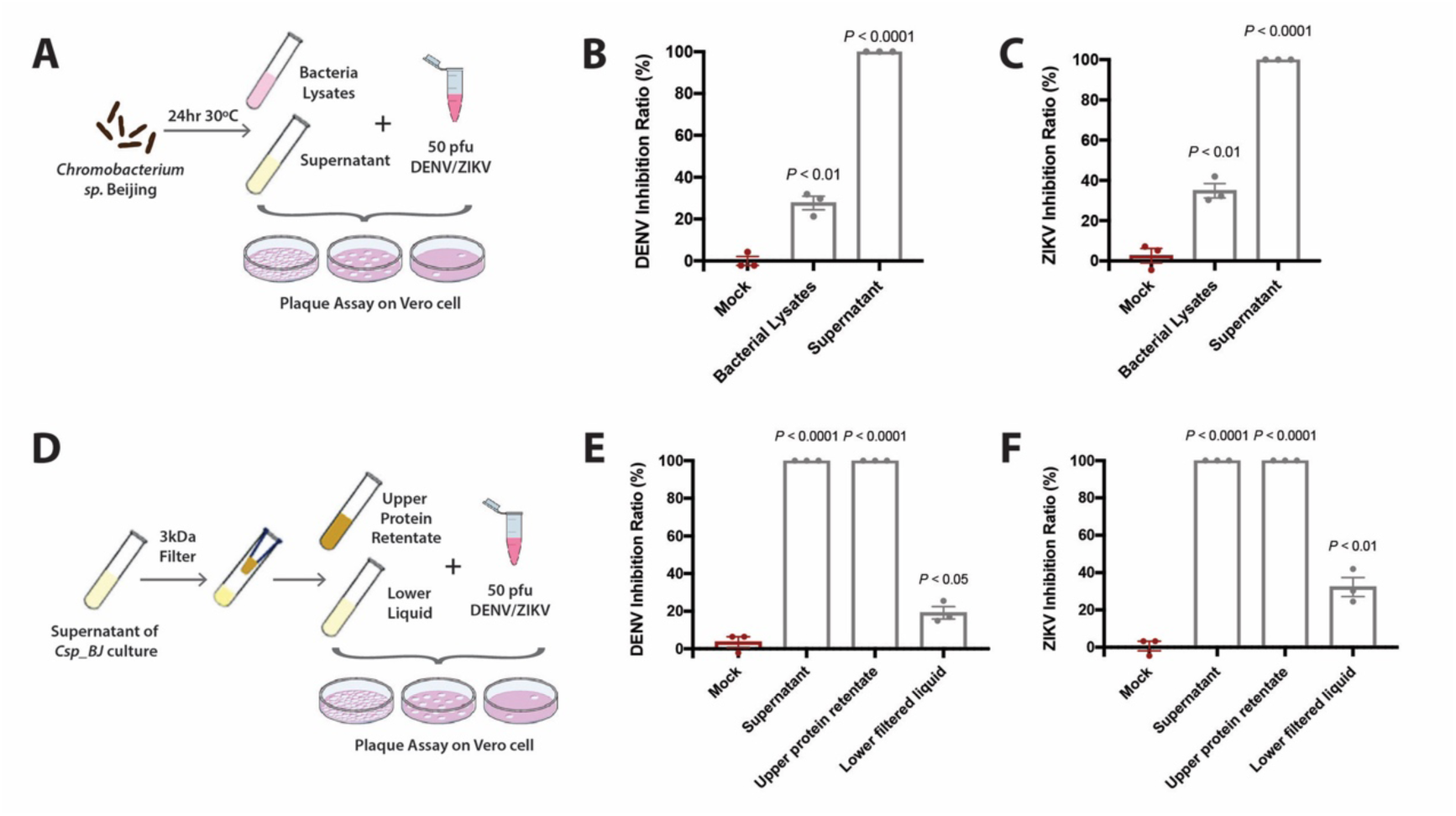
Secreted effector(s) from *Csp_BJ* inhibit DENV and ZIKV infection in Vero cells. (A-C) Secreted factor(s) from *Csp_BJ* inhibited DENV and ZIKV infection in Vero cells. (A) Schematic representation of the study design. A *Csp_BJ* suspension was separated into bacterial cells and culture supernatant by centrifugation. Either the bacterial lysates or the culture supernatant (50% v/v) mixed with 50 pfu of DENV or ZIKV in VP-SFM medium (50% v/v) were incubated for 1 hr before being used for infection of Vero cell monolayers. Vero cells infected with fresh LB broth mixed with virus-containing medium served as a negative control. (B, C) Inhibition rate in the presence of cell lysates or culture supernatant of *Csp_BJ* following infection by DENV (B) or ZIKV (C) in Vero cells, as determined by plaque formation assay. (D–F) Proteins in the supernatant inhibited DENV and ZIKV infection in Vero cells. (D) Schematic representation of the study design. Either the retentate (proteins) or the filtrate (small molecules and short peptides) (50% v/v) was mixed with 50 pfu of DENV or ZIKV in VP-SFM medium (50% v/v) and incubated for 1 hr before inoculation into Vero cell monolayers. Vero cells infected with fresh LB broth or culture supernatant mixed with virus-containing medium served as negative or positive controls, respectively. (E, F) The inhibition rate in Vero cells was determined by plaque formation assay. (B, C, E, F) Significance was determined by unpaired t-tests. Data are presented as the mean ± SEM.

**Figure 2.**
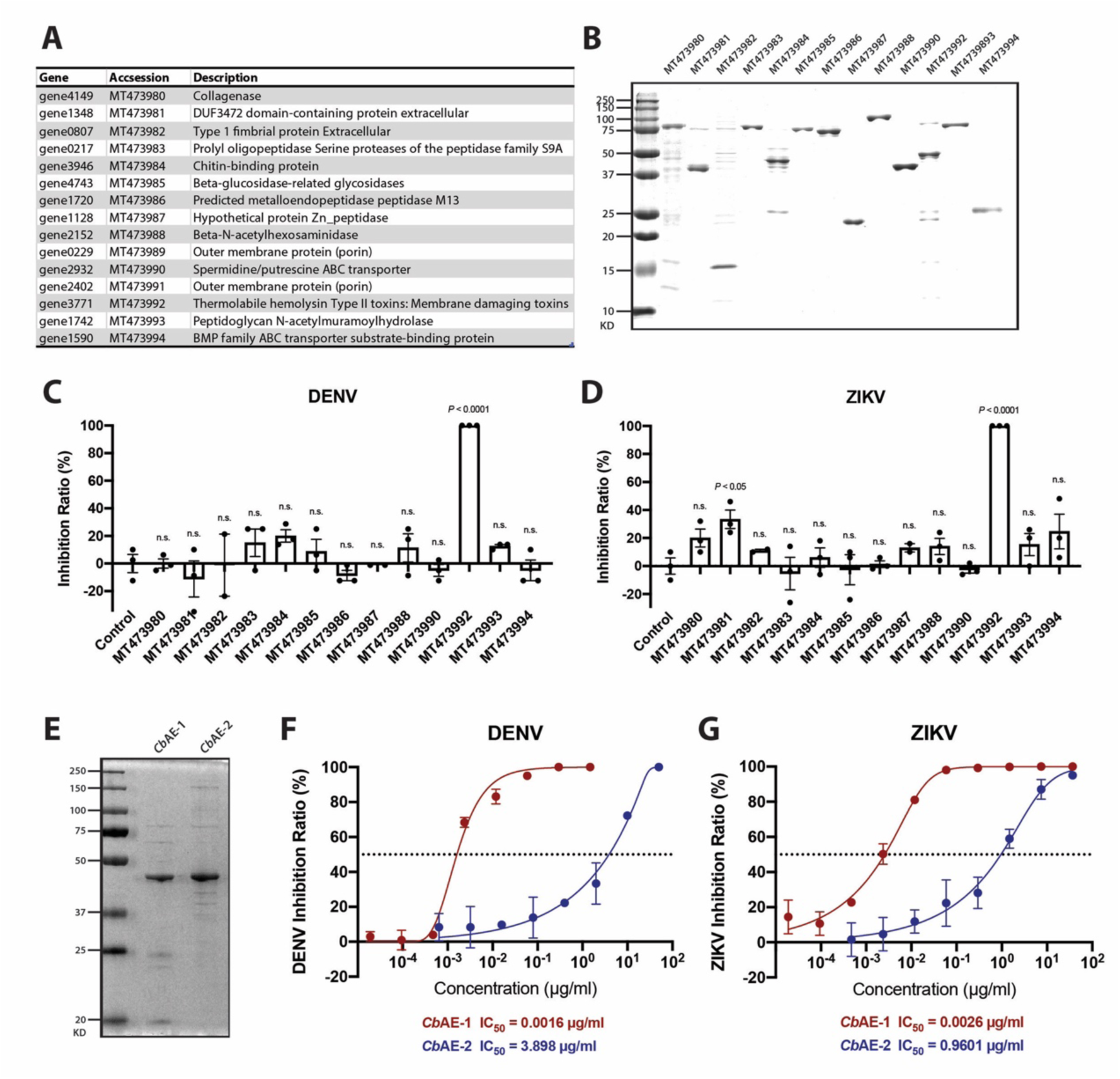
Identification of *Csp_BJ* secreted antiviral effector(s) against DENV or ZIKV. (A) The protein description was obtained from the UniProt and NCBI databases. (B) The identified *Csp_BJ* proteins were expressed and purified from *E. coli* cells. (C, D) A total of 1 µg of purified recombinant protein was mixed with 50 pfu of DENV (C) or ZIKV (D) in VP-SFM medium and incubated for 1 hr before infecting Vero cell monolayers. (E-G) *Csp_BJ* inhibited DENV and ZIKV infections via its secreted proteins *Cb*AE-1 and *Cb*AE-2. (E) The identified *Csp_BJ* secreted proteins *Cb*AE-1 and *Cb*AE-2 were expressed and purified from *E. coli* cells. (F, G) Inhibition curves of *Cb*AE-1 and *Cb*AE-2 against DENV (F) and ZIKV (G). Serial concentrations of *Cb*AE-1 or *Cb*AE-2 were mixed with 50 pfu of DENV or ZIKV in VP-SFM medium to perform standard plaque reduction neutralization tests (PRNTs). (C, D) Significance was determined using unpaired t-tests. Data are presented as the mean ± SEM.

We next investigated the mechanisms by which these bacterial effectors resist viral infection. According to sequence analysis, *Cb*AEs contain a conserved lipase domain. Lipases are a group of enzymes that catalyse the hydrolysis of the ester bond of glycerides into fatty acids and glycerol (Casas-Godoy et al., 2018). We therefore assessed whether *Cb*AEs have lipase activity. In a plate degradation assay, both *Cb*AEs directly digested egg yolk lipoproteins and formed lytic halos whose diameters correlated with lipase activity (Figure 3A). The sequence GDSL is the core motif of lipase activity (Casas-Godoy et al., 2018). Consistently, a S187G mutation in this motif of *Cb*AE-1 fully disrupted its lipase activity (Figure 3A), validating *Cb*AEs as secreted lipases of *Csp_BJ*.

**Figure 3.**
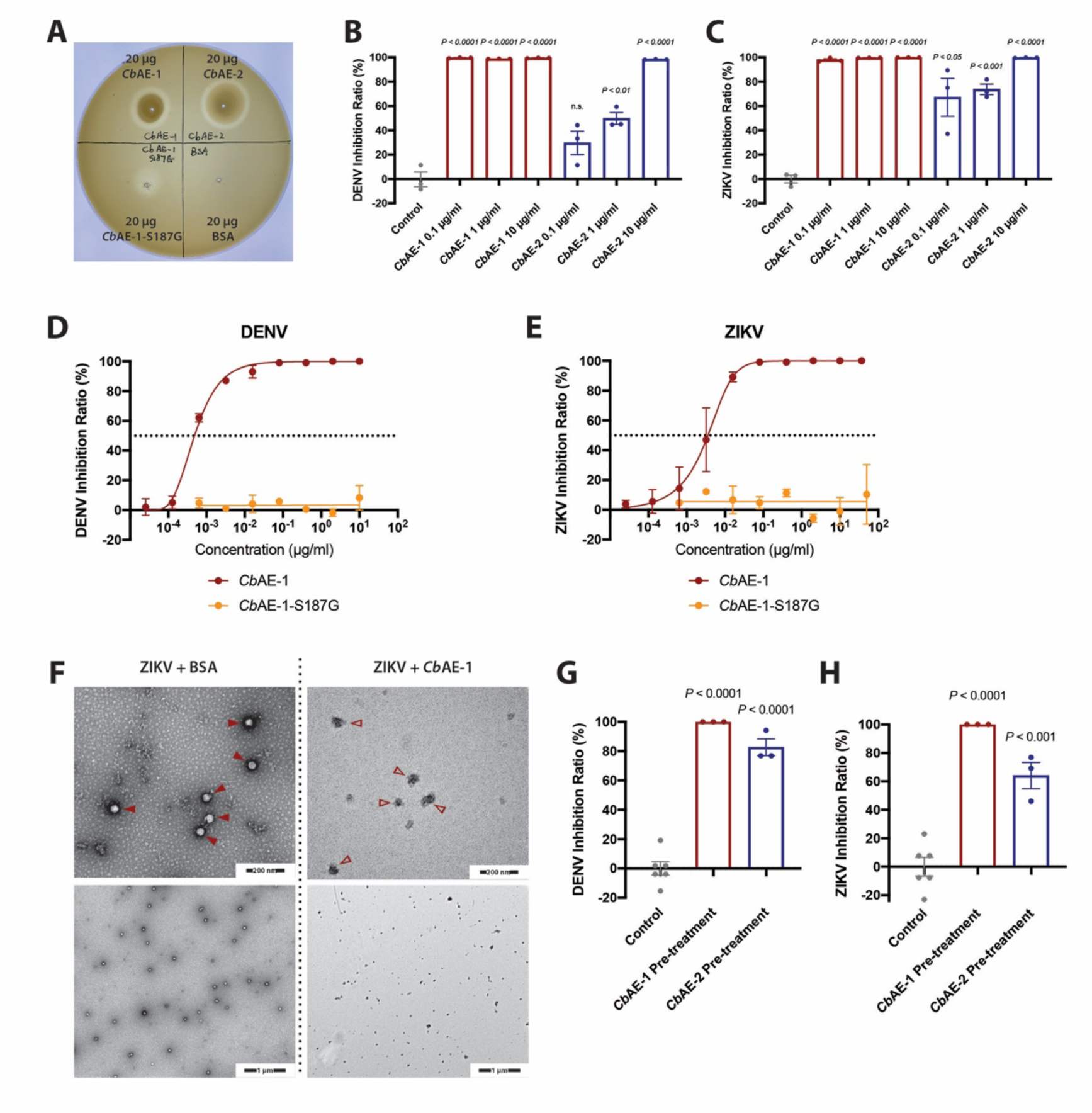
The virucidal activity of *Cb*AEs is mediated by enzymatic degradation of the viral lipidic envelope. (A) Lipase enzymatic activity of *Cb*AE-1, *Cb*AE-2 and *Cb*AE-1-S187G measured via an egg yolk agar plate assay. (B, C) Analysis of the exposure of DENV (B) or ZIKV (C) genomic RNA. DENV or ZIKV was first treated with serial concentrations of *Cb*AE-1, *Cb*AE-2 or BSA and then with RNase-A. Viral RNA degradation was evaluated by RT-qPCR. (D, E) The S187G mutant of *Cb*AE-1 fully lost its ability to suppress DENV (D) and ZIKV (E) infection: inhibition curves of *Cb*AE-1 and *Cb*AE-1-S187G against DENV (D) or ZIKV(E). Serial concentrations of *Cb*AE-1 or *Cb*AE-1-S187G were mixed with 50 pfu of DENV or ZIKV in VP-SFM medium to perform standard plaque reduction neutralization tests (PRNTs). (F) Representative negative stained transmission electron microscopy images of ZIKV particles treated with 10 µg/ml BSA (arrow head) and those treated with 10 µg/ml *Cb*AE-1 (empty arrow head); high magnification: 120,000×, low magnification: 30,000×. (G, H) Rate of DENV (G) or ZIKV (H) replication inhibition following exposure to *Cb*AEs before viral infection of Vero cell monolayers. The viral genome was quantified by RT-qPCR. (B, C, G, H) Significance was determined using unpaired t-tests. Data are presented as the mean ± SEM.

Given their lipase activity, we hypothesized that *Cb*AEs might use their enzymatic activity to degrade viral lipid envelope, which may result in exposure of the viral RNA (Muller et al., 2014). To address this hypothesis, serial concentrations of *Cb*AEs were incubated with 1×10^4^ pfu of DENV or ZIKV for 1 hr at 37°C. The mixture was then treated with RNase A to evaluate the degradation of exposed viral genomic RNA. Compared to mock treatments in which the viruses were incubated with BSA, a significant reduction in viral RNA was recorded by RT-qPCR when the viruses were treated with *Cb*AEs (Figure 3B and 3C), indicating that the lipase activity of *Cb*AEs directly disrupted the virion structure, thus resulting in viral genome release. Consistent with these results, the S187G mutant of *Cb*AE-1 that had no lipase activity completely failed to suppress both DENV and ZIKV infection (Figure 3D and 3E), further indicating that the virucidal activity of *Cb*AEs is lipase-dependent. To validate that *Cb*AEs disrupt viral lipidic membranes, we incubated *Cb*AE-1 with purified ZIKV virions and processed the samples for transmission electron microscopy (Figure 3F). The ZIKV particle typically has a diameter of 50–60 nm (Hasan et al., 2018) and, in consistent with that, we observed intact viral particles in our control sample treated with 10 µg/ml BSA (Figure 3F). However, upon treatment with 10 µg/ml *Cb*AE-1 in the same buffer as BSA solution, the integrity of ZIKV particles was fully disrupted (Figure 3F). This is in agreement with our previous finding that treatment of *Cb*AEs resulted in viral genome release. Since *Cb*AEs blocked viral infection in the extracellular milieu, we further assessed whether treatment of *Cb*AEs prior to viral inoculation could effectively block viral infection. We pre-treated Vero cells with 10 µg/ml of purified *Cb*AEs. Subsequently, 0.1 MOI of DENV or ZIKV was used to challenge the *Cb*AEs-treated cells. Pre-incubation with *Cb*AE-1 fully blocked the infectivity of both flaviviruses, whereas treatment with *Cb*AE-2 exhibited 60%-80% inhibition (Figure 3F and 3G).

Since the *Cb*AEs showed a direct catalytic action on the viral lipid bilayer, we assessed their virucidal activity against other enveloped viruses. SARS-CoV-2 is a newly emerging coronavirus that causes the severe acute respiratory disease COVID-19. To date, no specific therapeutics are available against SARS-CoV-2 infection. We therefore assessed the virucidal effect of the *Cb*AEs on SARS-CoV-2. In contrast to the large difference in virucidal activity against flavivirus infection, both *Cb*AEs presented a similar antiviral effect against a SARS-CoV-2 pseudovirus in HEK-293T-ACE2 cells (Figure 4A). Consistently, infection with low-passage SARS-CoV-2 in Vero cells was also significantly suppressed by treatment with the *Cb*AEs (Figure 4B). Additionally, both *Cb*AEs showed broad-spectrum antiviral effects on HIV-1 pseudoviruses (Figure 4C, 4D and 4E), as well as HSV-1 (Figure 4F). Notably, compared to a HIV-neutralizing antibody (N6) with a near-pan neutralization breadth (Huang et al., 2016), *Cb*AEs were very potent with a much lower IC_50_ for 3 HIV pseudovirus strains (Figure 4C, 4D and 4E). Nonetheless, it is puzzling that the *Cb*AEs did not show any effect against infection with influenza A virus (IAV) (Figure 4G).

**Figure 4.**
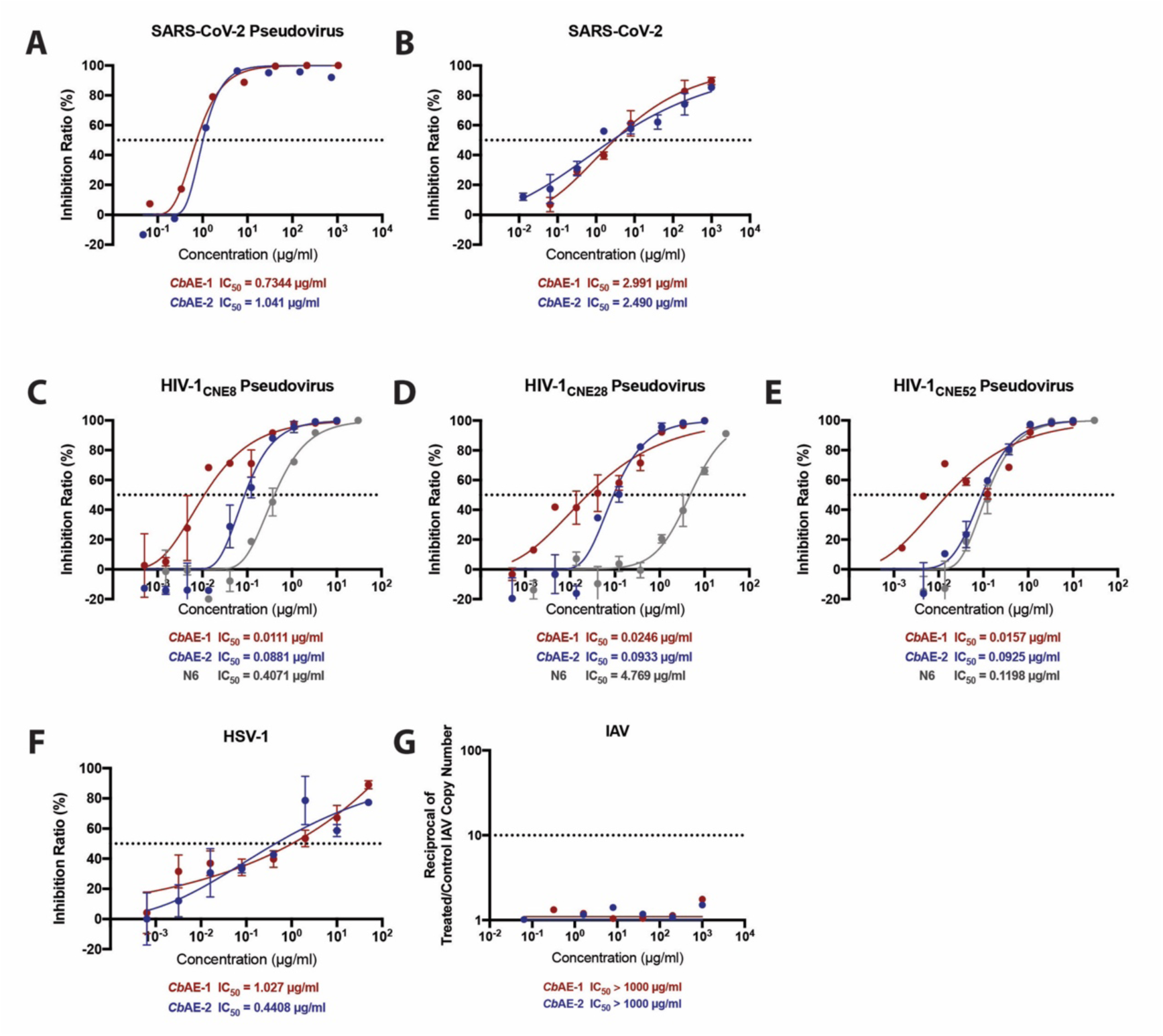
The virucidal activity of *Cb*AEs against 4 enveloped viruses. (A-G) Inhibition curves of *Cb*AE-1 and *Cb*AE-2 against SARS-CoV-2 pseudovirus (A), SARS-CoV-2 (B), HIV-1 pseudovirus CNE8 strain (C), HIV-1 pseudovirus CNE28 strain (D), HIV-1 pseudovirus CNE52 strain (E), HSV-1 (F) or IAV (G). Standard plaque reduction neutralization tests (PRNTs) (B, F), luciferase-based neutralization assays (A, C-E) or a RT-qPCR-based neutralization assay (G) was performed.

The lipidic envelope of a viral particle is generally derived from the plasma membrane or the endoplasmic reticulum (ER) membrane of a host cell. Thus, *Cb*AEs can also act on host cell membranes and their cytotoxicity to host cells may be a major concern for developing a *Cb*AE-based broad-spectrum antiviral drug. Generally, *Cb*AE-1 showed much higher toxicity than *Cb*AE-2 on both Vero cells (Supplementary Figure 3A) and A549 human adenocarcinomic epithelial cells (Supplementary Figure 3B). The half-cytotoxicity concentration (CC_50_) of *Cb*AE-2 was >1,000 µg/ml in the tested mammalian cells (Supplementary Figure 3A and 3B). The selectivity index (SI = CC_50_/IC_50_) of *Cb*AE-2 was >1000 for all the viruses/cells tested; while the SI of *Cb*AE-1 ranges from 200 to 400,000. We next assessed the toxicity of the *Cb*AEs in ICR mice. The half-lethal dose (LD_50_) was either 91.05 µg/kg for *Cb*AE-1 or 4000 µg/kg for *Cb*AE-2 upon tail vein injection (Supplementary Figure 3C). Nonetheless, the mice were much tolerant with a high dose of *Cb*AEs via intranasal inoculation. The animals intranasally treated with a high dose of *Cb*AEs all survived and maintained normal body weight (Supplementary Figure 3D). Altogether, these data suggest that *Cb*AE-2 is much safer than *Cb*AE-1 to hosts, and thus could be a broad antiviral drug candidate.

In this study, we identified two virucidal effectors with lipase activity, *Cb*AE-1 and *Cb*AE-2, from the *Chromobacterium sp. Csp_BJ*. Both *Cb*AEs showed a potent virucidal activity against a variety of enveloped viruses including DENV, ZIKV, SARS-CoV-2, HIV-1, and HSV-1. Notably, neither *Cb*AEs exhibited any effect against IAV. The toxicity assessment showed that *Cb*AE-2 was much safer than *Cb*AE-1 to both human cells and mice. Indeed, accumulating evidence indicates that certain lipases present a potent antiviral activity. Either lipoprotein lipase or hepatic triglyceride lipase impaired hepatitis C virus (HCV) infection in human Huh7.5 cells through degrading virus-associated lipoproteins (Shimizu et al., 2010). A secreted phospholipase A2 (PLA2) isolated from *Naja mossambica* snake venom showed a potent virucidal activity against HCV, DENV, and Japanese encephalitis virus (JEV); while the protein did not exhibit significant antiviral activity against Sindbis virus (SINV), IAV, Middle East respiratory syndrome coronavirus (MERS-CoV) or HSV-1 (Chen et al., 2017). A secreted human PLA2 has been shown to neutralize HIV-1 by degrading the viral membrane (Kim et al., 2007) or by blocking viral entry into host cells rather than a lipase-mediated virucidal effect (Fenard et al., 1999), suggesting diverse virucidal mechanisms by PLA2. *Cb*AEs may inactivate viruses by their lipase activity that also damage cellular membranes; nonetheless, their specificity and affinity for the viral envelope could be significantly improved, thus further reducing their cytotoxicity and increasing virucidal efficacy. For example, the SARS-CoV-2 surface protein Spike binds to human ACE2 with high affinity (Wang et al., 2020), and thus soluble ACE2 could be engineered with *Cb*AEs to greatly enhance its affinity and specificity for the viral envelope. Given that human recombinant soluble ACE2 can also inhibit SARS-CoV-2 infection in organoids (Monteil et al., 2020), it is possible that a soluble ACE2-*Cb*AE construction may have improved efficacy in blocking SARS-CoV-2 infections.

Viruses are the major aetiological agents of acute severe human diseases that impose a tremendous burden on the global public health and economy (Ghosn et al., 2018; Girard et al., 2010; Wilder-Smith et al., 2017). Given the fact that development of virus-specific vaccines and antiviral drugs is usually lengthy, broad-spectrum antiviral drugs could be crucial to prevent wide spread of new viral disease in a timely manner (Schein, 2020). *Cb*AE-2, with a broad antiviral effect and low toxicity to hosts, could be a potential choice. Our study provides a future avenue for the development of broad-spectrum antiviral drugs that might reduce the clinical burden caused by emerging viral diseases.

## Acknowledgements

This work was funded by grants from the National Key Research and Development Plan of China (2016ZX10004001-008 and 2017YFC1201004), the Natural Science Foundation of China (81730063, 81961160737 and 31825001), and the Shenzhen San-Ming Project for prevention and research on vector-borne diseases (SZSM201611064). We thank the core facilities of the Center for Life Sciences and Center of Biomedical Analysis (Tsinghua University) for technical assistance.

## Materials and Methods

### Ethics statement

Human blood was collected from healthy donors who provided written informed consents. The collection of human blood samples and their use for mosquito feeding was approved by the local ethics committee at Tsinghua University.

### Mice, mosquitoes, cells, viruses and bacteria

Eight-week-old female ICR mice purchased from Vital River Laboratories in China were used for toxicity assay. The mice were maintained in a specific pathogen-free barrier facility at Tsinghua University. The animal protocol used in this study was approved by the Institutional Animal Care and Use Committee of Tsinghua University and performed in accordance with their guidelines. *Aedes aegypti* (the Rockefeller strain) was maintained on a sugar solution in a low-temperature, illuminated incubator (Model 818, Thermo Electron Corporation) at 28°C and 80% humidity, according to standard rearing procedures (Cheng et al., 2010). Vero cells, HEK-293T cells and A549 cells were maintained in Dulbecco’s modified Eagle’s medium (11965-092, Gibco) supplemented with 10% heat-inactivated foetal bovine serum (16000-044, Gibco) and 1% antibiotic-antimycotic (15240-062, Invitrogen) in a humidified 5% (V/V) CO_2_ incubator at 37°C. The Vero, HEK-293T and A549 cell lines were purchased from the ATCC (CCL-81, CRL-3216 and CCL-185, respectively). DENV-2 (New Guinea C strain), ZIKV (PRVABC59 strain), HSV-1, IAV (H1N1 PR8 strain) and SARS-CoV-2 were grown in Vero cells with VP-SFM medium (11681-020, Gibco). DENV, ZIKV, HSV and SARS-CoV-2 were titrated by a standard plaque formation assay on Vero cells (Bai et al., 2007). IAV were titrated using a standard 50% tissue culture infection dose (TCID_50_) assay (Teferedegne et al., 2013; Varada et al., 2013). All experiments involving infectious SARS-CoV-2 were performed in a biosafety level 3 (BSL3) containment laboratory. *Chromobacterium sp.* Beijing was grown in LB broth at 30°C for 24 hr at 250 rpm. Culture supernatants were obtained by centrifugation and filtering the supernatant through a 0.22 µm filter unit (SLGP033RS, Millipore).

### Mass spectrometry

*Chromobacterium sp.* Beijing was washed three times with PBS and suspended in 10 ml of VP-SFM medium. After incubation for 2 hr at 37°C, the bacteria were pelleted and removed by centrifugation and filtration using a 0.22 µm filter unit (SLGP033RS, Millipore). The protein component of the bacterial supernatant was concentrated using an Ultra-15 centrifugal filter concentrator (UFC900396, Amicon) and subjected to SDS-PAGE. Fresh VP-SFM medium served as a negative control. The whole gel lane was excised and analysed by liquid chromatography-mass spectrometry (LC-MS) at the Protein Chemistry Technology Core, Tsinghua University. The MS readouts were searched against the protein sequence database of *Chromobacterium spp.* at the UniProt Database using Mascot software. The secreted proteins with a score ≥ 800 were included in the subsequent investigation.

### Protein expression and purification

The genes identified from mass spectrometry were amplified from *Csp_BJ* cDNA and cloned into the pET-28a (+) expression vector. Recombinant proteins were induced to be expressed in the *E. coli* BL21 DE3 strain using 200 mM IPTG for protein expression for the soluble form and 500 mM IPTG for inclusion bodies. The proteins were induced by 200 mM IPTG overnight at 16°C to generate soluble forms and purified with TALON metal affinity resin (635501, Clontech). The proteins were eluted with 250 mM imidazole and subsequently dialyzed in PBS buffer (pH 7.4). The inclusion bodies were washed with lysis buffer (50 mM Tris, 150 mM NaCl, 5 mM CaCl_2_, 5% Triton X-100 and 1 mM DTT) 3 times and 2 M urea 1 time, dissolved in 8 M urea and dialyzed overnight in renaturation buffer. Endotoxin was removed (L00338, GenScript) before the protein concentration was measured using a Bradford assay (500-0006, Bio-Rad), and the protein purity was checked with SDS-PAGE.

### Plaque reduction neutralization tests (PRNTs)

Vero cells were seeded at ∼4×10^5^ cells per well in 6-well plates and then incubated at 37°C overnight before reaching 80–90% confluence. DENV, ZIKV, HSV-1 and SARS-CoV-2 virus stocks were diluted to 50 plaque-forming units (pfu) per ml and incubated untreated or with a serial dilution of the *Cb*AEs in five-fold steps at 37°C for 1 hr before being added onto Vero cell monolayers for 2 hr of infection. Cell monolayers were washed once with PBS and covered with 1% agarose overlay DMEM with 2% FBS. After 4-5 dpi (DENV, ZIKV and HSV-1) or 2-3 dpi (SARS-CoV-2), Vero cell monolayers were fixed and stained with 0.8% crystal violet, and the number of pfu per ml was determined. The concentration of each protein necessary to inhibit virus infection by 50% (IC_50_) was calculated by comparison with the untreated cells using the dose-response-inhibition model in GraphPad Prism 8.0 (GraphPad Software, USA).

### Pre-treatment virus inhibition assays

For pre-infection treatment, Vero cells were seeded in 24-well plates and allowed to form monolayers. Ten micrograms/ml *Cb*AE-1 or *Cb*AE-2 was added to Vero cell monolayers at 37°C for 1 hr, and then DENV or ZIKV (0.1 MOI) was added and incubated for another hour at 37°C. After infection, cell monolayers were washed once with PBS buffer, fresh VP-SFM medium was added, and the cells were incubated at 37°C for 48 hr before the supernatant was collected for RT-qPCR quantitation of the viral genome.

### Viral genome quantitation by RT-qPCR

Total RNA was isolated either from homogenized mosquitoes or infected cell supernatant using a Multisource RNA Miniprep Kit (AP-MN-MS-RNA-250, Axygen) and reverse transcribed to cDNAs using an iScript cDNA Synthesis Kit (1708890, Bio-Rad). Viral genomes were quantified by qPCR using iTaq Universal SYBR Green Supermix (1725121, Bio-Rad). RT-qPCR was performed on a Bio-Rad CFX-96 Touch Real-Time Detection System. Primer sequences are shown in Supplementary Table 1.

### Lipase activity assay

The lipase activity of *Cb*AE-1, *Cb*AE-2 and *Cb*AE-1-S187G was measured with a plate assay as previously described (Liu et al., 1996). Briefly, 20 µg of *Cb*AE-1, *Cb*AE-2 or *Cb*AE-1-S187G was spot inoculated onto a 2% agar plate with 1% egg yolk and incubated for 24 hr at 37°C. Phospholipase activity was indicated by the diameter of the lytic halo around each well.

### Viral RNA exposure assay

DENV (1×10^4^ pfu) or ZIKV (1×10^4^ pfu) was incubated with different concentrations of the *Cb*AEs or PBS in a total volume of 1 ml for 1 hr at 37°C. Then, the mixtures were treated with 1 µl of RNase A (GE101-01, Transgen) and incubated for 1 hr at 37°C. Viral RNA was extracted, and RNA degradation was evaluated by RT-qPCR as mentioned above.

### Transmission electron microscopy

ZIKV viral particles were purified as described previously (Tan and Lok, 2014). Briefly, virus stocks were pelleted in 8% w/v PEG 8000 at 10,000 ×g for 1 hr, then purified by 24% w/v sucrose cushion for 2 hr at 175, 000 ×g (Beckman SW41 Ti rotor) and separated in potassium tartrate-glycerol gradient 10–35% for 2 hrs at 175,000 ×g (Beckman SW41 Ti rotor). Purified viral particles were suspended in 10 µg/ml BSA solution or 10 µg/ml *Cb*AE-1 solution and incubated for 1 hr at room temperature. The samples were then applied to a carbon grid, washed 3 times with water and negatively stained with 1% w/v uranyl acetate. The images were acquired in a Hitachi H-7650B TEM microscope at 80.0 kV.

### RT-qPCR-based neutralization assay

The *in vitro* antiviral efficacy of the *Cb*AEs on IAV was tested in A549 cells. Briefly, A549 cells were seeded into a 24-well plate and incubated at 37°C for 20-24 h. IAV (100 TCID_50_) were incubated untreated or with a serial dilution of the *Cb*AEs in five-fold steps at 37°C for 1 hr before being added onto A549 cell monolayers to allow infection to proceed for 2 hr. Then, the virus-protein mixture was removed, and the cells were further cultured with fresh VP-SFM medium. At 24 hr p.i., relative viral RNA copy numbers in the infected cells were quantified by RT-qPCR assays with specific primers. The neutralization titre was defined as the concentration of each protein necessary to inhibit the PCR signal by 90% (i.e., below the threshold of 10% of the mean value observed in virus control wells).

### HIV-1 and SARS-CoV-2 pseudovirus production and neutralization assay

HIV-1 pseudoviruses were generated by co-transfecting HEK-293T cells with Env expression vectors and the pNL4-3R-Eluciferase viral backbone plasmid, and a neutralization assay was performed as described previously (Zhou et al., 2019). Briefly, pseudovirus titres were measured by luciferase activity in relative light units (RLUs) (Bright-Glo Luciferase Assay System, Promega Biosciences, California, USA). Neutralization assays were performed by adding 100 TCID_50_ (median tissue culture infectious dose) of pseudovirus into 10 serial 1:3 dilutions of *Cb*AE-1 or *Cb*AE-2 starting from 50 µg/ml, following incubation at 37°C for 1 hr and addition of GhostX4/R5 cells. Neutralizing activity was measured by the reduction in luciferase activity compared to that in the controls. The fifty percent inhibitory concentration (IC_50_) was calculated using the dose-response-inhibition model with the 5-parameter Hill slope equation in GraphPad Prism 8.0 (GraphPad Software, USA). Vesicular stomatitis virus G protein (VSV-G) pseudotyped lentiviruses expressing human ACE2 were produced by transient co-transfection of pMD2G (Addgene #12259) and psPAX2 (Addgene #12260) plasmids and the transfer vector pLVX-ACE2Flag-IRES-Puro with VigoFect DNA transfection reagent (Vigorous) into HEK-293T cells to generate the HEK-293T-ACE2 cells for SARS-CoV-2 pseudovirus infection. SARS-CoV-2 pseudoviruses were purchased from GenScript, and neutralization activity was measured using the HEK-293T-ACE2 cell line with the same procedures as mentioned above.

### Membrane blood feeding

Fresh human blood from healthy donors was placed in heparin-coated tubes (367884, BD Vacutainer) and centrifuged at 1,000 ×g and 4°C for 10 min to separate plasma from blood cells. The plasma was heat-inactivated at 55°C for 60 min. The separated blood cells were washed three times with PBS to remove the anticoagulant. The blood cells were then resuspended in heat-inactivated plasma. Bacterial suspension (0.6 OD) was mixed with viruses and treated blood for mosquito oral feeding via a Hemotek system (6W1, Hemotek). Fully engorged female mosquitoes were transferred into new containers and maintained under standard conditions for an additional 8 days. The mosquitoes were subsequently euthanized for further analysis. The mosquitoes used in this experiment were previously antibiotic treated. Briefly, mosquitoes were provided with cotton balls moistened with a 10% sucrose solution including 20 units of penicillin and 20 mg of streptomycin per ml (15070-063, Thermo Fisher Scientific) for 5 days to remove gut bacteria. The mosquitoes were starved for 24 hr to allow the antibiotics to be metabolized prior to *in vitro* membrane blood feeding. Removal of gut bacteria was confirmed by a colony-forming unit assay.

### Cytotoxicity assay

The cytotoxicity of the *Cb*AEs was evaluated in Vero cells and A549 cells. Cell viability was measured by the MTT [3-(4,5-dimethylthiazol-2-yl)-2,5-diphenyl tetrazolium bromide] (M8180, Solarbio) method. Confluent cell monolayers contained in 48-well plates were exposed to different concentrations of the *Cb*AEs for 24 hr at 37°C. Then, a final concentration of 0.5 mg/ml MTT was added to each well. After 4 hr of incubation at 37°C, the supernatant was removed, and 250 µl of dimethyl sulfoxide (DMSO) was added to each well to solubilize the formazan crystals. After shaking for 10 min, absorbance was measured at 490 nm. The concentration of each protein necessary to reduce cell viability by 50% (CC_50_) was calculated by comparison with the untreated cells using a sigmoidal nonlinear regression function to fit the dose-response curve in GraphPad Prism 8.0 (GraphPad Software, USA).

### Toxicity study in ICR mice

ICR mice were used to test the *in vivo* safety of the *Cb*AEs. A number of 108 ICR 8-week old female mice were divided into 27 groups (n=4) at random. *Cb*AE-1 or *Cb*AE-2 was dissolved in PBS and administered either intravenously once at doses of 64, 320, 1600, 4000 and 8000 µg/kg or intranasally once at doses of 64, 320, 1600 and 8000 µg/kg. PBS and corresponding concentrations of BSA served as negative controls. The general behaviour, signs of toxicity, body weights and mortality of the mice were recorded after the administration of the *Cb*AEs. The half-lethal dose (LD50) was calculated using a sigmoidal nonlinear regression function to fit the dose-response curve in GraphPad Prism 8.0 (GraphPad Software, USA).

### Quantification and Statistical Analysis

Animals were randomly allocated into different groups. Mosquitoes that died before sample collection were excluded from the analysis. The investigators were not blinded to the allocation during the experiments or to the outcome assessment. No statistical methods were used to predetermine the sample size. Descriptive statistics are provided in the figure legends. All analyses were performed using GraphPad Prism statistical software.

## Supplementary Information

**Supplementary Figure 1.**
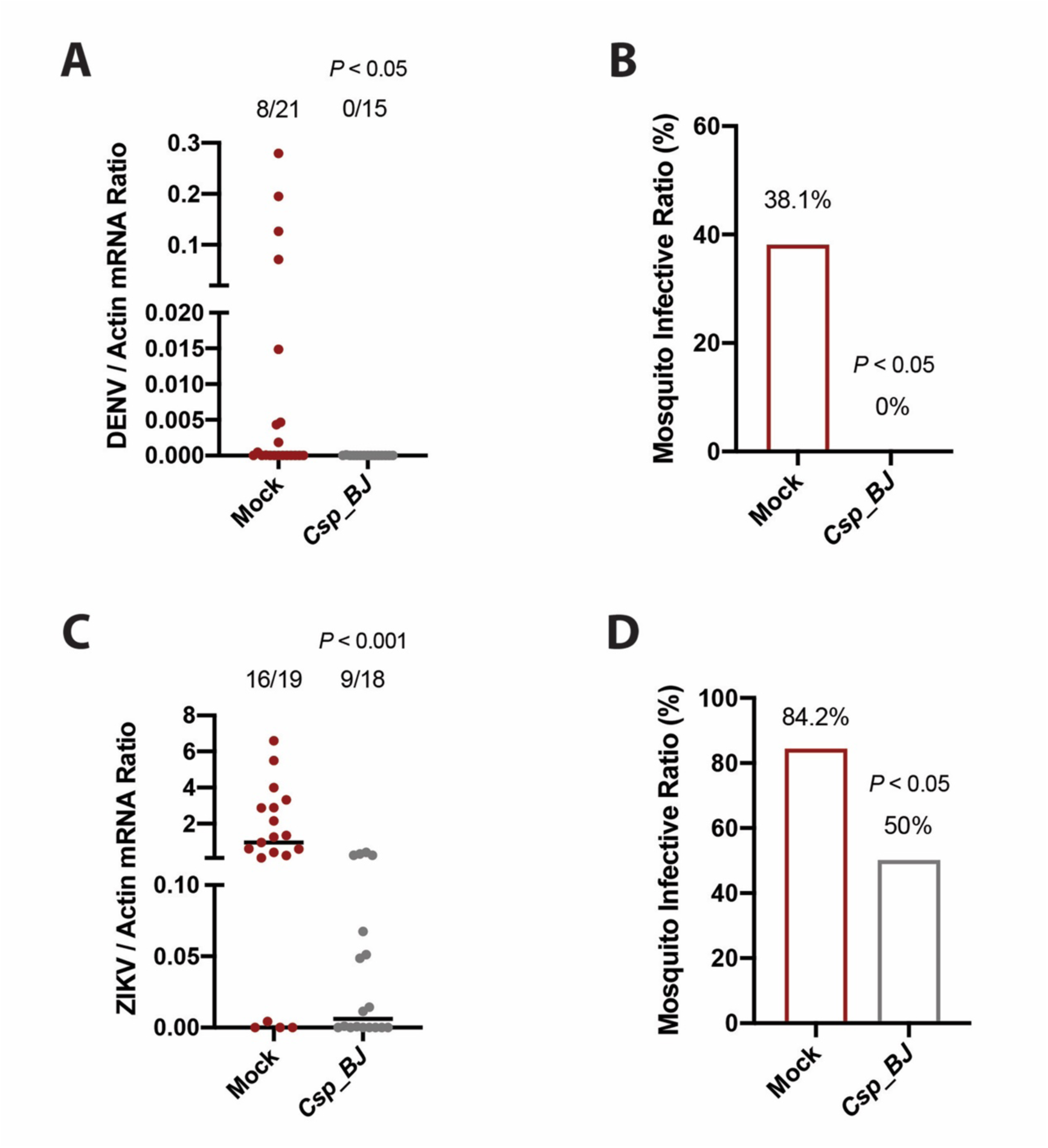
*Csp_BJ* inhibits DENV or ZIKV infection in *Aedes aegypti*. (A-D) Supplementation of *Csp_BJ* inhibits DENV (A, B) and ZIKV (C, D) infection of *A. aegypti*. A mixture containing human blood (25% v/v), *Csp_BJ* bacterial suspension (25% v/v), and supernatant from DENV- or ZIKV-infected Vero cells (50% v/v) was used to feed antibiotic-treated *A. aegypti* Rockefeller strain via an *in vitro* blood feeding system. Mosquito infectivity was determined by RT-qPCR at 8 days post blood meal. The final DENV or ZIKV titre was 1 × 10^5^ pfu/mL for oral infection. (A, C) The number of infected mosquitoes relative to total mosquitoes is shown at the top of each column. A nonparametric *Mann-Whitney* test was used for the statistical analysis. (B, D) Differences in the infectivity ratio were compared using *Fisher’s* exact test.

**Supplementary Figure 2.**
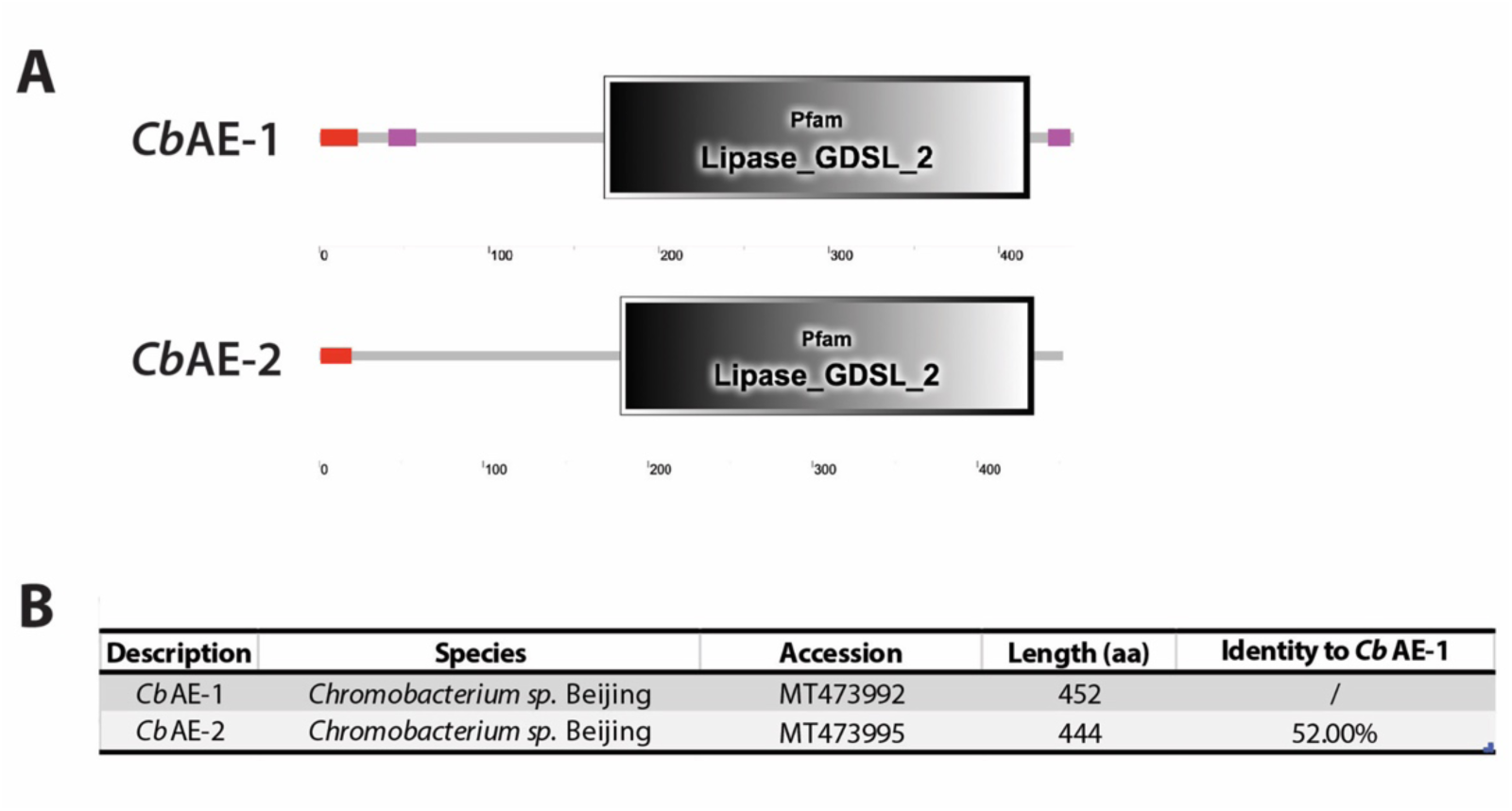
Sequence comparison of *Cb*AE-1 and its homologue in *Csp_BJ*. (A) Conserved domains of *Cb*AE-1 and *Cb*AE-2 protein sequences was analysed using a Simple Modular Architecture Research Tool (SMART) (Letunic and Bork, 2018; Letunic et al., 2015). (B) Sequence comparison of *Cb*AE-1 and *Cb*AE-2 was performed using Basic Local Alignment Search Tool (BLAST) on NCBI website with the program “Needleman-Wunsch alignment of two sequences” (Altschul et al., 1997).

**Supplementary Figure 3.**
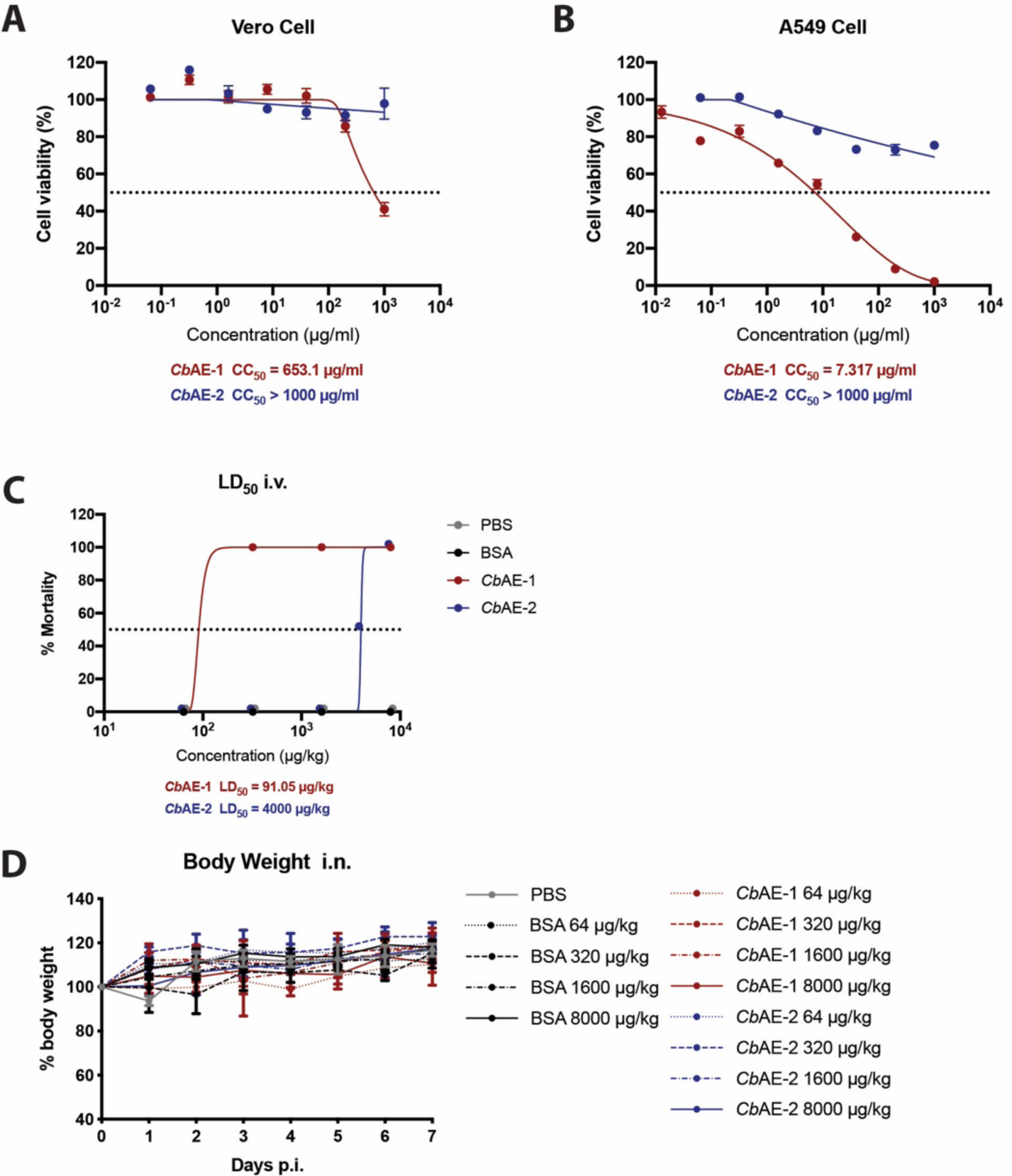
Toxicity evaluation of the *Cb*AEs in Vero cells, A549 cells and ICR mice. (A, B) Cytotoxicity of *Cb*AEs to Vero cells (A) or A549 cells (B) was measured by MTT assays. (C-D) Toxicity assay of *Cb*AEs in ICR mice. (C) Mortality rate of acute intravenous (i.v.) administration of the *Cb*AEs in ICR mice (n=4 per group). (D) Body weight during the 7-day monitoring period in ICR mice subjected to acute intranasal (i.n.) administration of the *Cb*AEs (n=4 per group).

**Supplementary Table 1.**
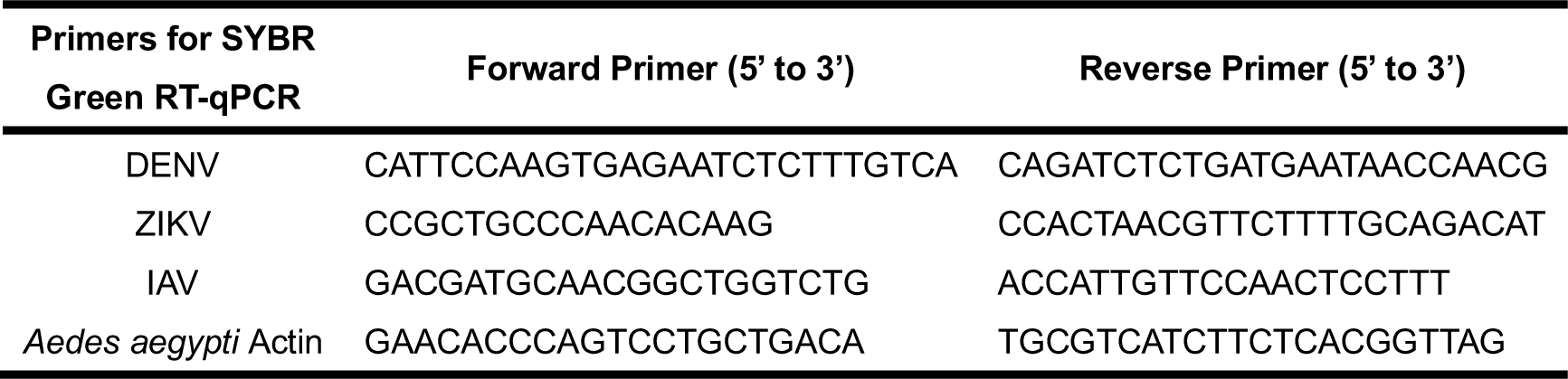
Primers for SYBR Green RT-qPCR.

## Notes

### Competing Interest Statement

The authors have declared no competing interest.

### Summary of Updates

Supplementary Figure 3 revised.

